# “Fire! Do Not Fire!”: A new paradigm testing how autonomous systems affect agency and moral decision-making

**DOI:** 10.1101/2023.12.19.572326

**Authors:** Adriana Salatino, Arthur Prével, Émilie Caspar, Salvatore Lo Bue

## Abstract

Autonomous systems have pervaded many aspects of human activities. However, research suggest that the interaction with those machines may influence human decision-making processes. These effects raise ethical concerns in moral situations. We created an ad hoc setup to investigate the effects of system autonomy on moral decision-making and human agency in a trolley-like dilemma. In a battlefield simulation, participants had to decide whether to initiate an attack depending on conflicting moral values. Our results suggest that our paradigm is suitable for future research aimed at understanding the effects of system autonomy on moral decision -making and human agency.

## 1. Introduction

In the last decades, the use of autonomous systems has become increasingly widespread in various fields of human activity. From driving (Ayoub et al., 2019; Chan, 2017) to aviation (Anderson et al., 2018; Chialastri, 2012; Valdés et al., 2018), medicine (Kawamoto et al., 2005; Sutton et al., 2020), and military defense and security (Mayer, 2015), there is almost no area where these technologies are not massively deployed. As these technologies become more widespread, the inevitable trend toward even more machine autonomy will lead to profound changes in the role played by humans during the execution of tasks. From workers in industry, agriculture, and transportation, to consumers in their daily lives, advances observed in technologies show that humans are rapidly moving from the position of direct tasks executors (with the help of mechanistic machines) to supervising tasks performed directly by intelligent machines with high level of autonomy.

The main reason for the massive deployment of autonomous systems resides in the many benefits these systems offer to users. Several laboratory experiments have shown that the introduction of some level of autonomy in tasks can have substantial effects in terms of users’ decision-making and performance. For example, studies have shown that autonomous systems can help people in detecting task-relevant cues and ignoring irrelevant cues in the environment (Chavaillaz et al., 2018; Goh et al., 2005), improve the decisions made by human subjects in complex situations and reduce the number of errors they make (MacMillan et al., 1997; Rovira et al., 2007; Sarter & Schroeder, 2001),or even reduce the time to make correct decisions (Chavaillaz et al., 2018). In addition, the introduction of autonomy has been shown to reduce users’ mental workload and thus increase their ability to monitor multiple tasks simultaneously (e.g., Chen & Barnes, 2012; Wright et al., 2018). Laboratory experiments have also shown that the effects produced by the interaction with autonomous systems on human decision-making and performance depend on several factors (Parasuraman & Riley, 1997; Mosier & Manzey, 2019). In particular, autonomous systems operating at the decision stage of a task or with a high level of autonomy are generally associated with the largest benefits in terms of human performance (e.g., Endsley & Kaber, 1999; Manzey et al., 2012; Rovira et al., 2007). Based on these results, one could easily conclude that more autonomy is always better for the users, whether they are pilots flying on a plane, physicians analyzing the test results from a patient, or military drone operators deployed in a war zone.

However, the involvement of autonomous systems technologies into human activities has not been systematically associated with positive effects. Several studies have shown that their use can also have significant negative effects. Parasuraman et al. (1993) provided a classic demonstration thereof. They reported low level of detection of automation failures with highly reliable systems, an effect they called automation complacency (also named automation overreliance). Other examples of negative effects are loss of situational awareness (Endsley, 2017), skill decay (Haslbeck & Hoemann, 2016; Volz & Dorneich, 2020), performance decrement in return-to-manual control (Endsley & Kiris, 1995), or increased workload with too many autonomous systems to monitor (Wang et al., 2009). Interestingly, the detrimental effects caused by the cooperation with autonomous systems seem to be directly related with the stage and level of autonomy of those systems. For example, Rovira et al. (2007) reported lower rates of correct decisions with highly reliable systems with high levels of autonomy (i.e., decision systems) compared to systems with lower levels of autonomy (i.e., information systems). Thus, while more autonomy seems to be clearly beneficial when the system’s recommendations are correct, the negative effects seem also to be more pronounced for higher levels of autonomy in machines that are, most of the time, imperfect.

In this context, another important aspect of human-autonomous systems interaction that has received increasing attention in recent years is the impact of autonomy on human agency (Berberian, 2012, 2019, Coyle et al., 2012; Zanatto et al., 2021). The sense of agency (SoA), defined as the *feeling of causing changes in the external world by controlling one’s own voluntary actions* (Jeannerod, 2003; Haggard, 2017; Burin et al., 2017; Pyasik, Salatino et al., 2019), is recognized as an important aspect of human consciousness. Because SoA enables us to perceive ourselves as causal agents, it is the basis for intentional behavior (Haggard & Tsakiris, 2009), and is closely related to moral responsibility (Moretto et al., 2011; Caspar, Christensen, Cleeremans, & Haggard, 2016).

The recent interest in this topic has been triggered by the possibility offered by the “Intentional Binding” effect to implicitly measure the SoA. The Intentional Binding is a phenomenon by which the perceived time between an action and its outcomes is modulated by the intentionality of that action. Time appears compressed in situations where the person is active, while time appears stretched in situations where the person is passive (Haggard et al., 2002). Measuring the SoA by using this effect usually consists of asking subjects to estimate the time interval between an action they perform and the consequences of that action. Numerous studies have now shown that the time estimation between action and outcome is a valid implicit measure of SoA (e.g., Christensen et al., 2019; Imaizumi & Tanno, 2019; Malik & Obhi, 2019; Haggard, Clark, and Kalogeras, 2002; Moore and Obhi, 2012) and is preferable to a subjective measurement of responsibility, which is usually obtained by a direct report of how people attribute the effects of their own actions (Saito et al., 2015), which is subject to social desirability and other biases (e.g., Blackwood et al., 2003; Wegner & Withley, 1999).

In one of the first studies investigating the effects of autonomy on SoA by using the Intentional Binding paradigm (Berberian et al., 2012), participants took part in a flight simulation and were assisted in their task by different levels of automation. Berberian and colleagues’ results showed a decrease in SoA with increasing levels of automation, suggesting that agency decreases with higher levels of automation. Further evidence using the same paradigm can be found in the study by Coyle and coworkers (2012), who investigated how assistance, in a machine -assisted point-and-click task, affects the user’s SoA. Their results suggest that, up to a certain point, the computer could assist users while also allowing them to maintain a sense of control over their actions and outcomes, hence of their agency. More recently, Zanatto et al. (2021) showed a similar negative impact of automation on SoA, and that the mental workload may also play a role in reducing agency. Taken together, these studies suggest that automation technology may affect the mechanism underlying human agency.

Hence, the evidence of negative effects on human decisions and performance, and the evidence of a decrease in the implicit and explicit SoA (Berberian et al., 2012; Coyle et al., 2012; Vantrepotte et al., 2022), that might result from the interaction with the autonomous systems, have serious performance and safety implications. Engineers working on the development of new forms of autonomous technologies should be aware of these effects and take them into account. Furthermore, these results have important implications when those systems are used in sensitive or moral domains such as in medicine or in the military, in which decisions of life and death have to be made.

To date, however, very little is known about how the interaction with an autonomous machine affects SoA and the decisions made by someone engaged in a moral scenario, and how this is influenced by the level of autonomy of the system. Indeed, it is possible that interacting with autonomous systems to make moral decisions negatively affects the moral and ethical decision-making process and the resulting actions, particularly in tasks and domains of moral value such as in the military context (Christensen et al., 2012; Cushman et al., 2013, 2017).

In recent years, research in the field of moral decision making and autonomy has focused mainly on the rules and/or algorithms that can be assigned to an autonomous system to perform ethical responses in moral situations (Arkin et al., 2011; Jiang et al., 2021). Surprisingly, until recently, little attention has been paid to understanding how a human agent’s ethical behavior in moral decision-making situations can be influenced by its interaction with an autonomous system (Köbis et al., 2021). The available data suggest a mixed picture of the effects of autonomous systems in social and moral decision-making situations. Indeed, while some recent evidence suggests that interaction with automation could lead to the promotion of prosocial behaviors (such as fairness and cooperation, see, e.g., de Melo et al., 2018, 2019), other studies have shown that it could also lead to unethical behaviors (Cohn et al., 2022; Leib et al., 2021). Concerning SoA, while some studies report that a moral context increases it (e.g., Moretto et al., 2011) and that a higher SoA is associated with higher prosocial decision-making (Caspar et al., 2022), it is not clear whether this is still true when decisions are made in collaboration with an autonomous intelligent machine.

Considering the lack of researchon how autonomous systemsimpact the SoA and decision-making in a moral context, and how this can be modulated by the level of autonomy of the system, in the present study, we aimed to build an ad hoc setup to investigate how the level of system autonomy affects SoA and the moral decision-making. To this end, we developed a task in which participants (military cadets) played the role of drone operators on a simulated battlefield and had to decide whether or not to initiate an attack, based on the presence of enemies and the risk that allies might also be harmed. Participants were exposed to three types of trials representing three types of uncertainty (*Moral Decision-Making* Trials, *No Risk* Trials, and *No Enemy* Trials) with three different levels of system autonomy, including no system assistance, information assistance (i.e., the system gives processed-information on the presence of enemies and the risk for allies), and decision assistance (i.e., the system provides a recommendation on the best decision to make). In our study, SoA is measured both at the implicit level, using the Intentional Binding paradigm, and at the explicit level through a subjective assessment of responsibility (using an ad-hoc scale). We also measured performance by using reaction time, the proportion of trials in which participants chose to attack, and the proportion of choices leading to the fewest ally losses (called *utilitarian choices* in our task).

The primary purpose of this research was to develop and test a new paradigm to investigate how the interaction with autonomous systems can affect the SoA and the decisions made by people when facing moral choices, and how the level of autonomy of the system influences this effect. Based on previous findings (Berberian at al., 2012, Coyle et al., 2012; Zanatto et al., 2021), we hypothesized that agency decreases with increasing levels of system autonomy, as indicated by a longer time estimation between action and outcome, and lower subjective judgments of responsibility at higher levels of system autonomy. We also hypothesized that if the SoA would be affected by the autonomous system, as well as the sense of responsibility associated with SoA, the moral decision-making would also be affected, with the number of attacks increasing as system autonomy increases (Caspar et al., 2018; Goh et al., 2005, Chavaillaz et al., 2018). In addition, we expected shorter reaction times and more utilitarian choices with higher levels of system autonomy. Crucially, by validating the new paradigm we propose with the present study, we hope to pave the way for new quantitative studies to understand how the interaction with autonomous systems affects agency and decision-making in a moral context. In turn, understanding these effects better could help in the development of safer and more efficient autonomous systems in the future.

## 2. Method

### 2.1. Participants

A total of 31 participants took part in the study (M_age_ = 22, SD = 2.28, Range = 19 – 36, 7 women, 24 men). Participants were cadets at the Royal Military Academy of Belgium and were recruited with the help of a Master student officer in the course of his thesis. Participants were in their third and fourth year of study, meaning they had notions of International Humanitarian Law, and thus about what is legally allowed and forbidden in the conduct of armed conflicts. The sample size was estimated using G*Power (Faul, Erdfelder, Lang, & Buchner, 2007), with a small-to-medium size effect f of 0.2, a threshold for significance α set at 0.05, and a power 1-β at .80. Based on these values, the estimated sample size was 28 participants. To compensate for potential data losses and exclusions, a total of 32 participants was targeted. However, considering the limited pool and recruitment period, 31 participants were finally included in this study. The study was performed in accordance with the principles expressed in the declaration of Helsinki and with the protocol of the local ethics review board at the Faculty of Psychology and Educational Sciences at Ghent University. Participants were informed of the general purpose and the duration of the experiment, and about their rights as participants in psychological research before giving their consent. Participation was voluntary and participants could withdraw their participation at any time without justification and without consequences. Written informed consent was obtained from each participant before the experiment.

### 2.2. Stimuli and procedure

All the material can be found on the Open Science Framework (Salatino et al., 2023). The experiment was programmed and presented using MATLAB 2020b and the Psychophysics Toolbox extension. The experiment was run on a laptop computer (display resolution: 2560*1600 pixels) and responses were collected via an AZERTY keyboard with left and right arrow keys serving as the response keys used in the experiment. The experiment was conducted in an experimental room in the department of Life Sciences at the Royal Military Academy. Participants were seated on a chair, in front of a table with the screen of the laptop being at approximately 40 cm from the participants. Each trial of the task consisted of a 50 by 50 grid with dark grey cells (RGB = [064, 064, 064]) presented on a black screen (RGB = [000, 000, 000]; see Fig. 1, below). Participants were instructed that the grid represented a radar display used to inform on the position of allies and enemies. Out of the 2500 cells, 100 were colored in light grey (RGB = [192, 192, 192]) and each light grey cell represented the position of a group of 10 allies. The number of allied forces (i.e., light grey cells) was kept constant across trials, but their position varied randomly from trials to trials. In addition, in 66% of trials one of the 2500 cells was colored in white (RGB = [255, 255, 255]) to represent the presence and the position of an enemy. Maximum one cell was colored in white during trials (i.e., maximum of one enemy). When an enemy (i.e., a white cell) was presented on a trial, participants were asked to choose between attacking the enemy or not by pushing the left (attack) and right (no attack) arrows of the keyboard (i.e., Action 1-A1, if they decide to attack, and Action 2-A2, if they decide to not attack). Participants were instructed that not pushing the “attack” button whenever an enemy was present would result in the death of five allies because of the continued hostile activities of that enemy. Instead, pushing the “attack” button whenever an enemy was present would result in the death of the enemy, but with sometimes a risk of collateral damages considering the position of the allies on the 50 by 50 grid radar display (see details in the next paragraph below). Collateral damage was likely, but not sure, when grey cells were separated from the enemy by less than five empty cells. When no enemy was shown on the grid, participants were asked to not push on the “attack” button.

**Fig. 1.**
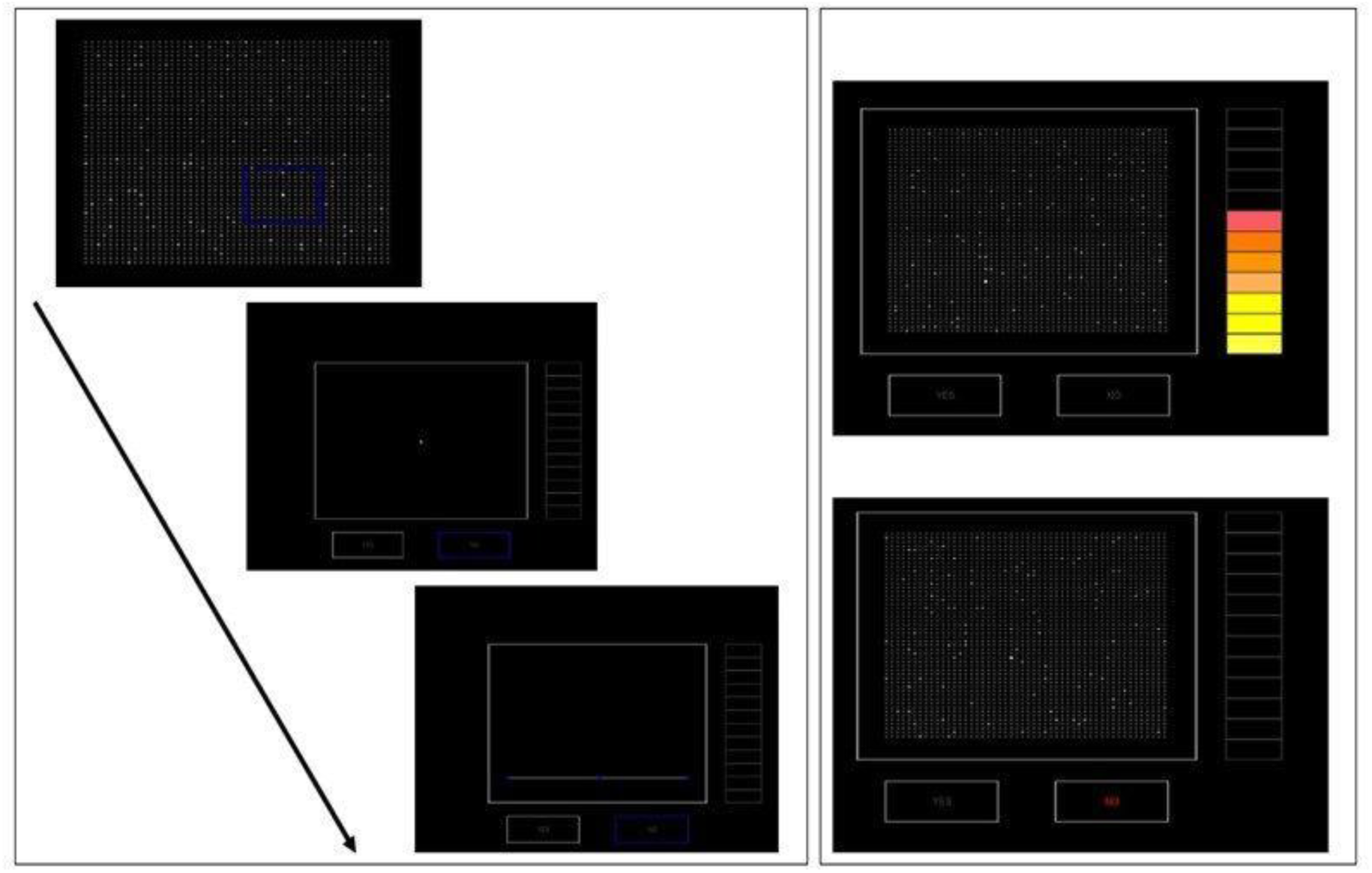
Experimental setup. Each trial of the task consisted of a 50 x 50 grid of dark grey cells on a black screen (**Left Panel**), representing a radar display informing of the position of allies and enemies. Of the 2500 cells, 100 were colored light grey and represented the position of a group of 10 allies, which varied in position from trial to trial. In 66% of the trials, one of the 2500 cells was coloured white to represent an enemy and participants were asked to choose whether or not to attack the enemy by pressing the left (attack) or right (no attack) arrow key on the keyboard. The grid was displayed for 15000ms or until the participant pressed one of the two response keys. After the response, a blackout screen was displayed for a random duration (200, 500 or 800ms) and a tone (200ms). Participants were asked to indicate the duration of the interval between their choice and the tone on a horizontal scale from 0 ms to 1000 ms. *Level 1* of system autonomy is shown in the right panel (**top**), where in addition to the basic visual assistance of *Level 0*, a scale has been displayed indicating the risk in terms of losses of allied forces in the event of an attack. In *Level 2* (**Lower Right Panel**), in addition to the visual assistance of *Level 0*, participants were assisted by a decision support system that gave a yes/no recommendation for the best decision to make.

During the experiment, participants were tested on three different types of trials, representing three levels of Uncertainty: *No Risk* trials, *No Enemy* trials, and *Moral Decision-Making (Moral DM)* trials. In the *No Risk trials*, an enemy was shown on the radar display and allies were separated from five cells of more from the enemy, meaning there were no risk of collateral damage. In the *No Enemy trials*, no enemy was shown on the radar display. Finally, in *Moral DM trials* an enemy was shown on the radar display and up to three groups of allies were on the first to the fifth range of cells next to the enemy. Here, participants received the instruction that during those trials there was a risk of collateral damages if they decided to attack. More exactly, participants were instructed that allies found on the 1 ^st^ range of cells next to the position of the enemy had a .50 probability of being killed if they decided to attack, while the probability of collateral damages decreased with the distance according to the following *range* – *probability* combinations: 2^nd^ range – .40, 3^rd^ range – .30, 4^th^ range – .20, and 5^th^ range – .10. The number of allies that could be killed during an attack varied from 10, 20, or 30 allies. Groups of allies were systematically on the same range of cells and *Moral DM* trials consisted in 15 different *number of allies* by *probabilities of collateral damages* combinations. Thus, we expected those trials to be morally challenging because participants were asked to choose between (1) pushing the attack button to neutralize the enemy, but with the risk of killing allies during the attack, or (2) not pushing the attack button and let the enemy kill 5 allies anyway.

The experiment consisted of three blocks, with each block including a specific level of Autonomy from intelligent system (*Level 0*, *Level 1*, *Level 2*). In *Level 0*, the block with the lowest level of assistance from intelligent system, participants received basic visual assistance to determine the position of allies relatively to the position of an enemy shown on the grid. During *No Risk* and *Moral DM* trials, by clicking with the help of the computer mouse on a white cell while the 50 by 50 grid was shown on the screen, the cells on the 6 ^th^ range next to the position of the enemy turned blue (RGB = [000, 000, 255]) until the participants made their decision. This visual information was designed to help the participants to detect how far allied forces were from the enemy’s position and to compute the risk of collateral damages if they decided to press the attack button. However, allies found within that area were at risk for collateral damages, with the risk depending on the range.

Then, in *Level 1* of Autonomy, in addition to the basic visual assistance found in *Level 0*, a 12 grade-scale was shown on the right part of the screen (see Fig. 1, right up) participants were instructed that this scale indicated the risk in terms of losses of allied forces if the attack button was pressed. The risk was computed based on the number of allies within the blue area (i.e., 10, 20, or 30) and the probability of collateral damages based on their position (i.e., .50, .40, .30, .20, or .10). The scale ranged from yellow (very low risk) to red (very high risk).

Finally, in *Level 2* of Autonomy, the visual assistance of *Level 0* was still included, but in addition participants were assisted by a decision-support system that on each trial made a yes/no recommendation on the best decision to make (see Fig. 1, right bottom). This recommendation was based in each trial on the choice associated with the lowest expected losses in terms of allied forces. When the expected losses were lower for pressing the attack button the ‘yes’ cue was highlighted, while the ‘no’ cue was highlighted when the expected losses were lower by pressing the no attack button (except during the trial where the number of allies was 10 and the probability was .50, in which expected losses were equal for both choice). Participants were not informed on the computation on which the recommendations were determined. Overall, each block consisted of 15 *No Risk* trials, 15 *No Enemy* trials, and 15 *Moral DM* trials.

The experimental setup of this task is shown in Fig. 1 (left): (1) First a loading bar was presented for delay chosen randomly between 1000ms and 2000ms to signal a new trial to the participants. (2) This was followed by a blackout screen for 500ms and then the presentation of the 50 by 50 grid, displayed for 15000ms or until the participant pressed one of the two response keys. (3) Participants were asked to confirm their choice by pressing the selected response key again, or they had the possibility to change their choice by pressing the other key. (4) Responses were followed by the presentation of blackout screen for a random duration of either 200, 500, or 800ms, and a tone (frequency: 400Hz) for 200ms. Finally, (5) participants were asked to report the duration of the interval between their confirmation choice and the tone on a horizontal scale ranging from 0 ms to 1000 ms. Trials were separated by an interval of 1000ms.

### 2.3. Measurements and analysis

We used five dependent variables in this study: Decision, Utilitarian Choice, Response Time, Agency, and Subjective Responsibility. Decision (A1) was expressed by the proportion of trials on which participants decided to attack. Utilitarian Choice (UC) was expressed by the proportion of choices implying the lowest expected losses (in percentage). Response Time (RT) was the mean response time (in seconds) on each trial. SoA was measured by Intentional Binding (IB, in milliseconds). IB was computed by subtracting each interval estimate from the mean actual response-tone interval (500ms) and averaged these scores for each Uncertainty X Autonomy condition. Each block of Autonomy ended with a subjective judgment of responsibility (SubjA), in which participants were asked to indicate how much they felt responsible of the decisions they made on a scale from −100 (not responsible at all) to 100 (entirely responsible)^1^.

Statistical analyses were performed using JASP version 0.17.2. We performed separate repeated-measures ANOVAs for A1, UC, RT, and IB with Uncertainty (*No Risk trials*, *No Enemy trials*, *Moral Decision-Making trials*) and Autonomy (*Level 0*, *Level 1*, *Level 2*) as within-subject factors. In addition, SubjA was compared by means of a repeated measures ANOVA with Autonomy as within-subject factors. For each dependent variable, only data of participants within +/− 2.5 SDs were considered. Greenhouse-Geisser correction was applied where sphericity was violated. We assessed Moral decision-making by several indicators.

The primary focus of our analysis concerned the presence of a main effect of Uncertainty on 1) A1, for which we expected a higher percentage of attacks during No Risk trials and a lower percentage during no Enemy trials, 2) UC, for which we expected an increased number in the No Risk and No Enemy (control) trials compared to the Moral Decision-Making trials, and 3) RT, with expected shorter response time in the No Risk and No Enemy trials compared to the Moral Decision-Making trials. These effects were expected to provide evidence of the moral conflict produced by the scenarios. Regarding IB, following the results of Moretto et al. (2011), we expected a main effect of Uncertainty with shorter time interval, indicating an increase of SoA, during *Moral Decision-Making* trials in comparison with the two control trials. We also expected a main effect of Autonomy on 1) UC, with an increased rate of UC with the level of autonomy of the task, 2) RT, congruent with the main effect of Uncertainty, 3) IB, with less IB, indicating a decrease of SoA, with increased level of autonomy in line with the conclusions of Berberian et al. (2012), and 4) SubjA, with lower SubjA with increased level of autonomy. These effects were expected to provide evidence of the effectiveness of the paradigm we developed. The threshold selected for significance was p < .05 with a two-tailed approach. Raw data, scripts, and processed data can be found on the Open Science Framework (Salatino et al., 2023).

## 3. Results

### 3.1. Analyses on A1 decisions

The analysis on A1 decisions (i.e. the proportion of attacks) (Fig. 2) revealed a main effect of Uncertainty (F (1.06, 27.72) = 926.70, p < .001, ηp2 = .97) and post hoc tests showed that all comparisons were significant (all ps < .001) with more a1 choices during *No Risk* trials (mean = 99.48, SEM = .20) in comparison with *Moral DM* trials (mean = 54.24, SEM = 1.77) and *No Enemy* trials (mean = .81, SEM = .52). However, the analysis revealed no significant effect of Autonomy (F (1.78, 46.39) = 2.80, p = .07, ηp2 = .09) on A1, as well as no significant interaction between Uncertainty and Autonomy (F (2.76, 71.98) = 1.17, p = .32, ηp2 = .04).

**Fig. 2.**
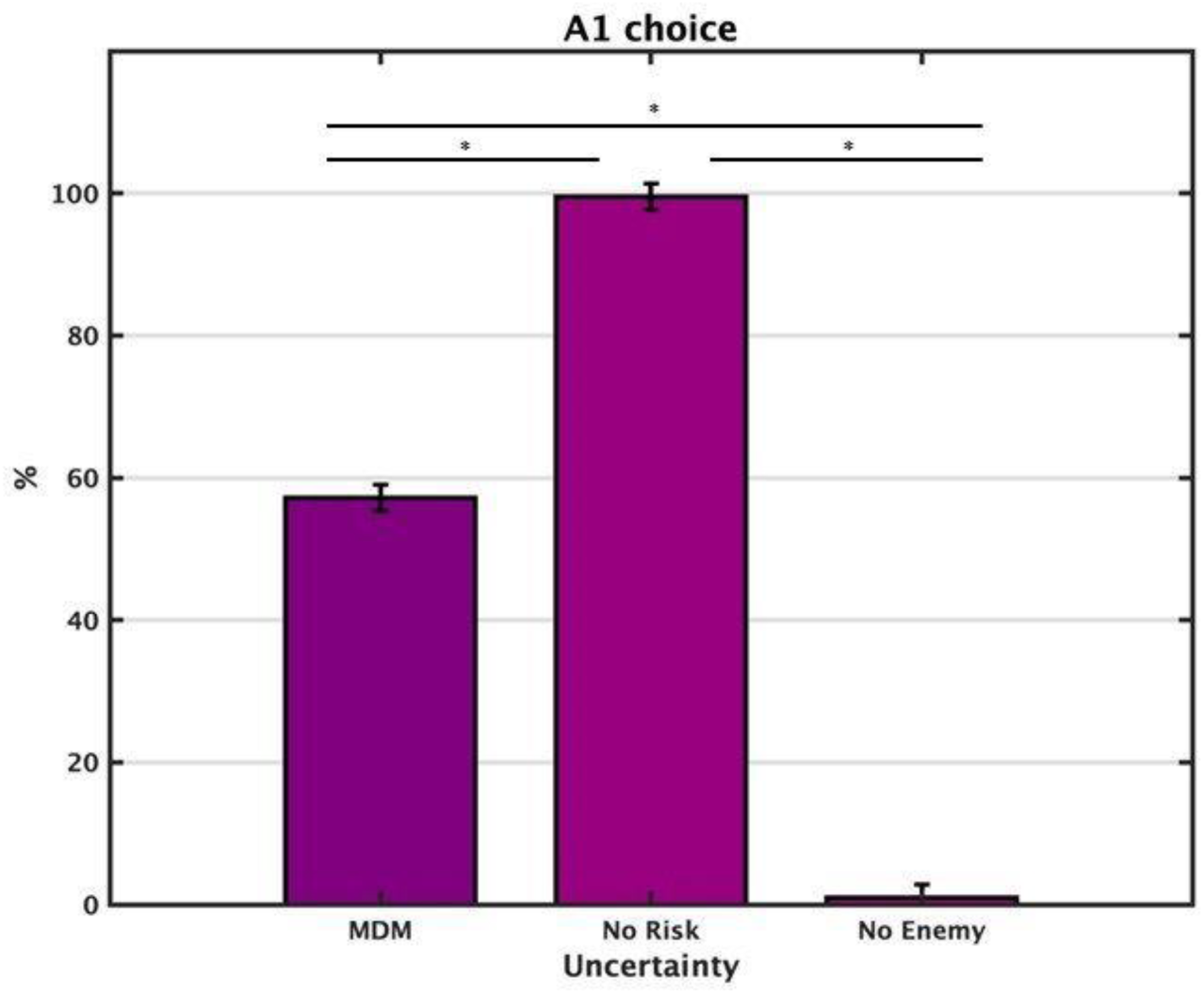
Proportion of A1 action (i.e. attacks performed). A significant increase of A1 choices during *No Risk* trials in comparison with *Moral Decision-Making* and *No Enemy* trials were found (all p < .001). * = significant.

### 3.2. Analyses on Utilitarian Choice (UC)

The analysis on UC (i.e., the choices implying the lowest expected losses, Fig. 3, Panel A) revealed a significant effect of Uncertainty (F (1.18, 31.96) = 95.76, p < .001, ηp2 = .78). Post hoc tests showed a significant difference between *Moral DM* trials (mean = 84.02, SEM = 1.09) and *No Risk* trials (mean = 99.30, SEM = 0.27), and between *Moral DM* and *No Enemy* trials (mean = 99.21, SEM =.50) (all ps < .001), with a reduced number of UC during *Moral DM* trials, but not between *No Risk* and *No Enemy* trials (p = 1.000). The analysis (Fig. 3, Panel B) also revealed a significant effect of Autonomy (F (1.76, 47.55) = 8.67, p < .001, ηp2 = .24). Post hoc tests showed no significant difference between *Level 0* (mean = 94.10, SEM = 1.07) and *Level 1* (mean = 92.59, SEM = 1.17) (p = .17) and between *Level 0* and *Level 2* (mean = 95.84, SEM = .88) (p = .089). A significant difference was found in the proportion of UC between *Level 1* and *Level 2* (p < .001) with more UC on *Level 2*. Finally, the analysis revealed no significant interaction between Uncertainty and Autonomy (F (2.36, 63.80) = 2.48, p = .08, ηp2 = .08).

**Fig. 3.**
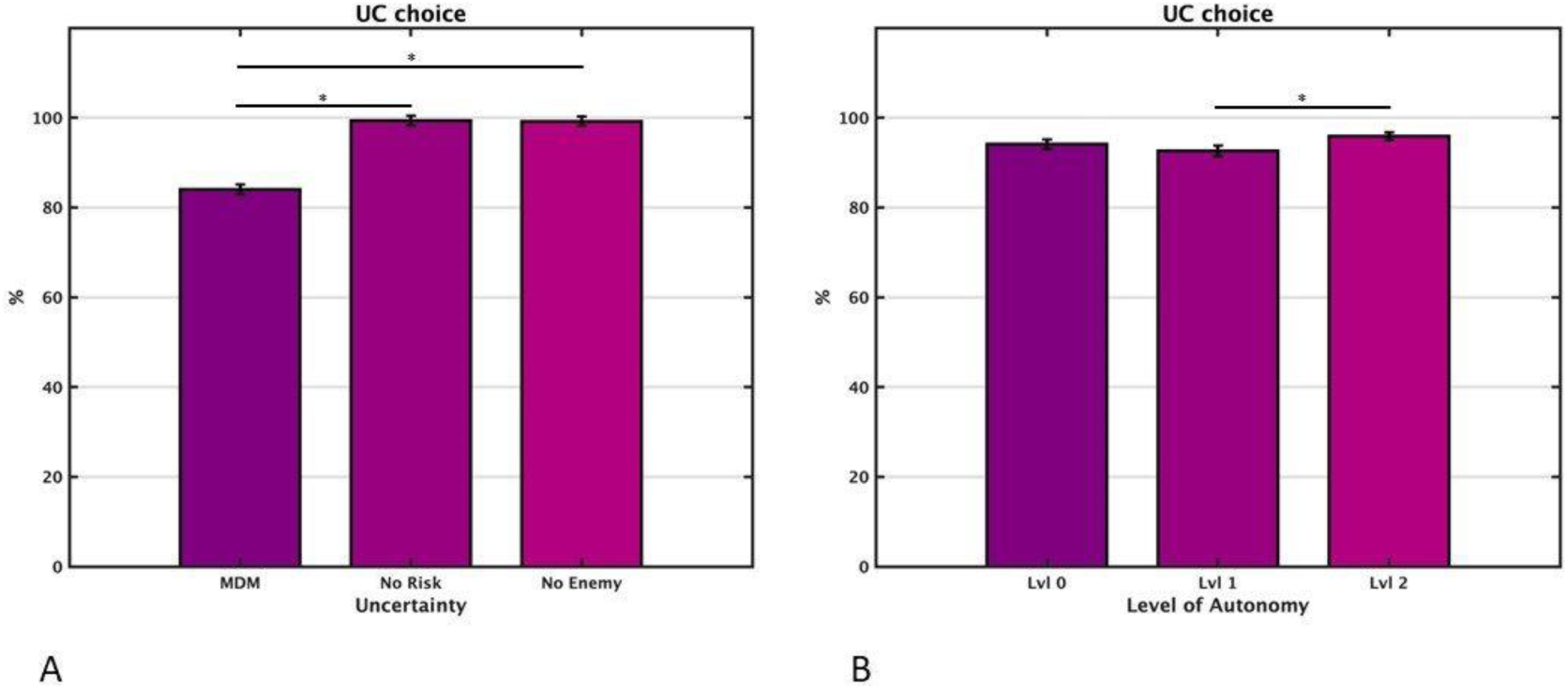
Proportion of UC (i.e. Utilitarian Choices). A significant difference was found in UC between *Moral Decision-Making* trials and *No Risk trials*, and between *Moral Decision-Making* and *No Enemy* trials (all p < .001), with a reduced number of UC during *Moral Decision-Making* trials, but not between *No Risk* and *No Enemy* trials (**Panel A**). A significant difference was found in the proportion of UC between *Level 1* and *Level 2* (p < .001) with more UC on *Level 2*. * = significant.

### 3.3. Analyses on Response Time (RT)

The analysis on RT (Fig. 4, Panel A) revealed a significant effect of Uncertainty (F (1.73, 46.95) = 68.76, p < .001, ηp2 = .71) with post hoc tests showing that all comparisons were significant (all ps ≤ 0.36), with longer RT during *Moral DM* trials (mean = 4.62, SEM = .20) than during *No Risk* (mean = 2.27, SEM = .12) and *No Enemy* trials (mean = 2.82, SEM = .14). In addition, the analysis revealed no significant effect of Autonomy (F (1.68, 45.36) = 0.667, p = .49, ηp2 = .02), but a significant interaction (Fig. 4, Panel B) between Uncertainty and Autonomy (F (3.43, 92.74) = 3.81, p = .009, ηp2 = .12), with a simple main effect of Autonomy in *No Enemy* trials (p = 0.03), but not in other Uncertainty conditions (all p > .37).

**Fig. 4.**
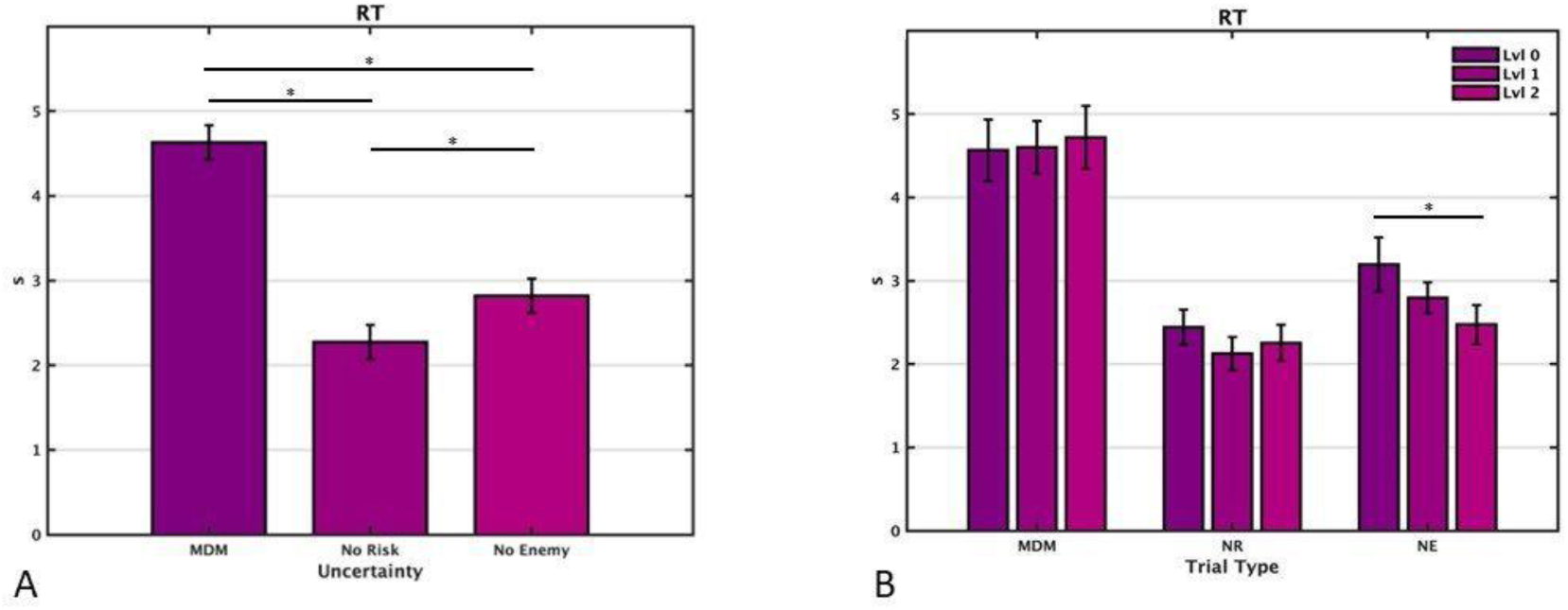
Response time (in seconds). A significant increase in the RT was found during *Moral Decision-Making trials* compared to the *No Risk* and *No Enemy trials* were found (all p < 0.36, **Panel A**). In addition, a significant (p = .009) interaction was found between Uncertainty and Autonomy (**Panel B**), with a simple main effect of Autonomy in *No Enemy trials* (p = 0.03). * = significant.

### 3.4. Analyses on Intentional Binding (IB)

The analysis on IB (Fig. 5) revealed a significant effect of Uncertainty (F (1.56, 46.87) = 8.12, p = .002, ηp2 = .21) with post hoc tests showing a significant difference between *Moral DM* trials (mean = 118.85, SEM = 13.58) and *No Risk* trials (mean = 147.04, SEM = 13.99) and between *Moral DM* and *No Enemy* trials (mean = 145.53, SEM = 13.69) (all p < 0.005), with shorter intervals reported in *No Risk* and *No Enemy trials* in comparison with *Moral Decision-Making* trials. No significant differences were found between *No Risk* and *No Enemy* trials (p = 1.000). Concerning Autonomy, the analysis revealed no significant effect (F (1.76, 52.82) = 0.67, p = .495, ηp2 = .022) and no significant interaction between Uncertainty and Autonomy (F (3.52, 105.86) = 1.29, p = .27, ηp2 = .04).

**Fig. 5.**
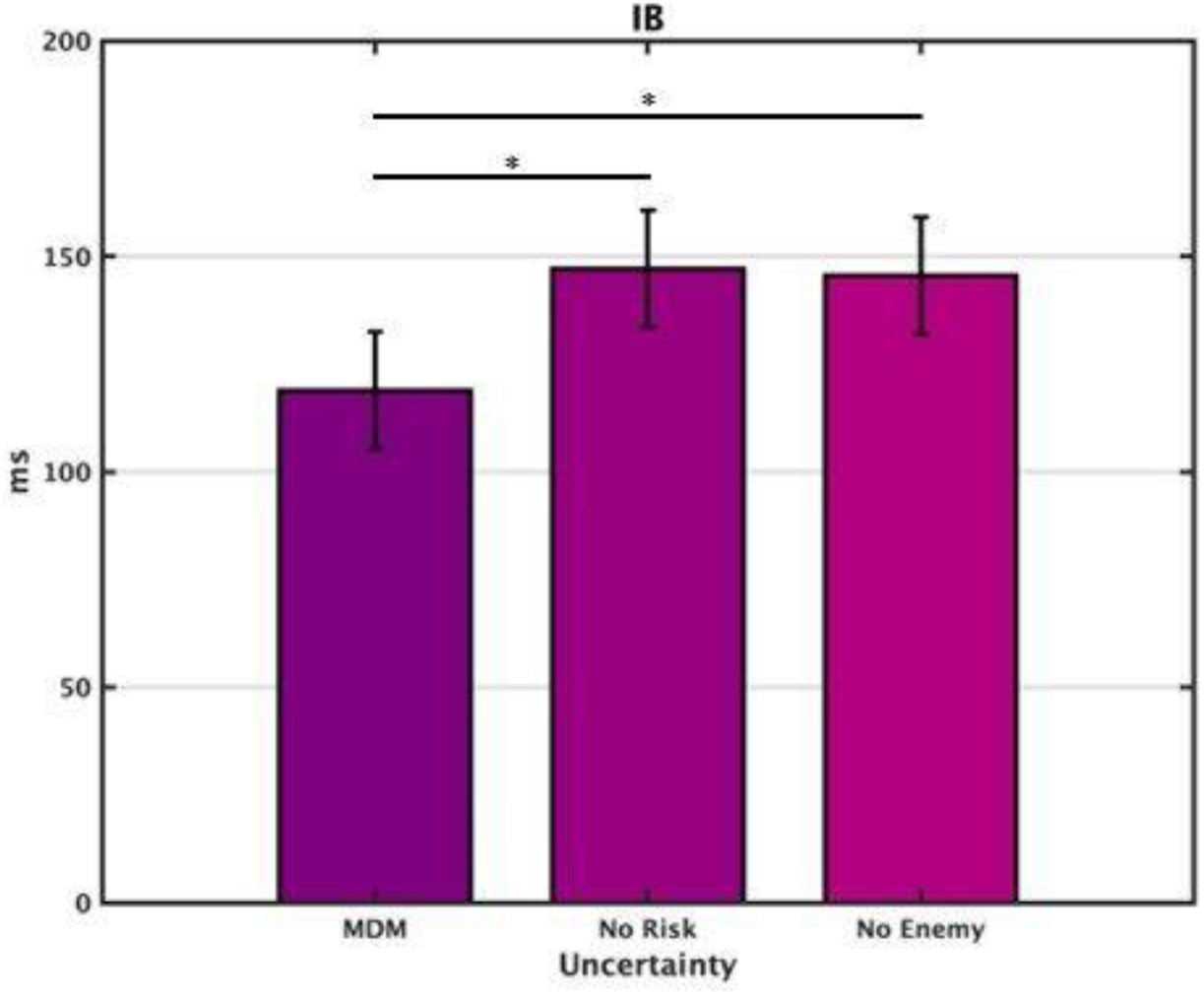
Intentional binding (IB). A significant difference in the IB was found between *Moral Decision-Making trials* and *No Risk trials*, and between *Moral Decision-Making trials* and *No Enemy trials* (all p < 0.005), with shorter intervals reported in *No Risk* and *No Enemy trials* compared to the *Moral Decision-Making trials*. * = significant.

### 3.5 Analyses of subjective judgment of responsibility (SubjAs)

The analysis on SubjAs (i.e., how much participants felt responsible of the decisions made, Fig. 6) revealed a significant effect of Autonomy (F (1.61, 45.15) = 15.72, p < .001, ηp2 = .36). Post hoc tests showed a significant difference between *Level 0* (mean = 84.13, SEM = 3.04) and *Level 1* (mean = 71.72, SEM = 4.60) (p = .006) and between *Level 0* and *Level 2* (mean = 58.62, SEM = 69) (p < .001), with larger subjective responsibility rating during *Level 0* in both cases, but not between *Level 1* and *Level 2* (p = .065).

**Fig. 6.**
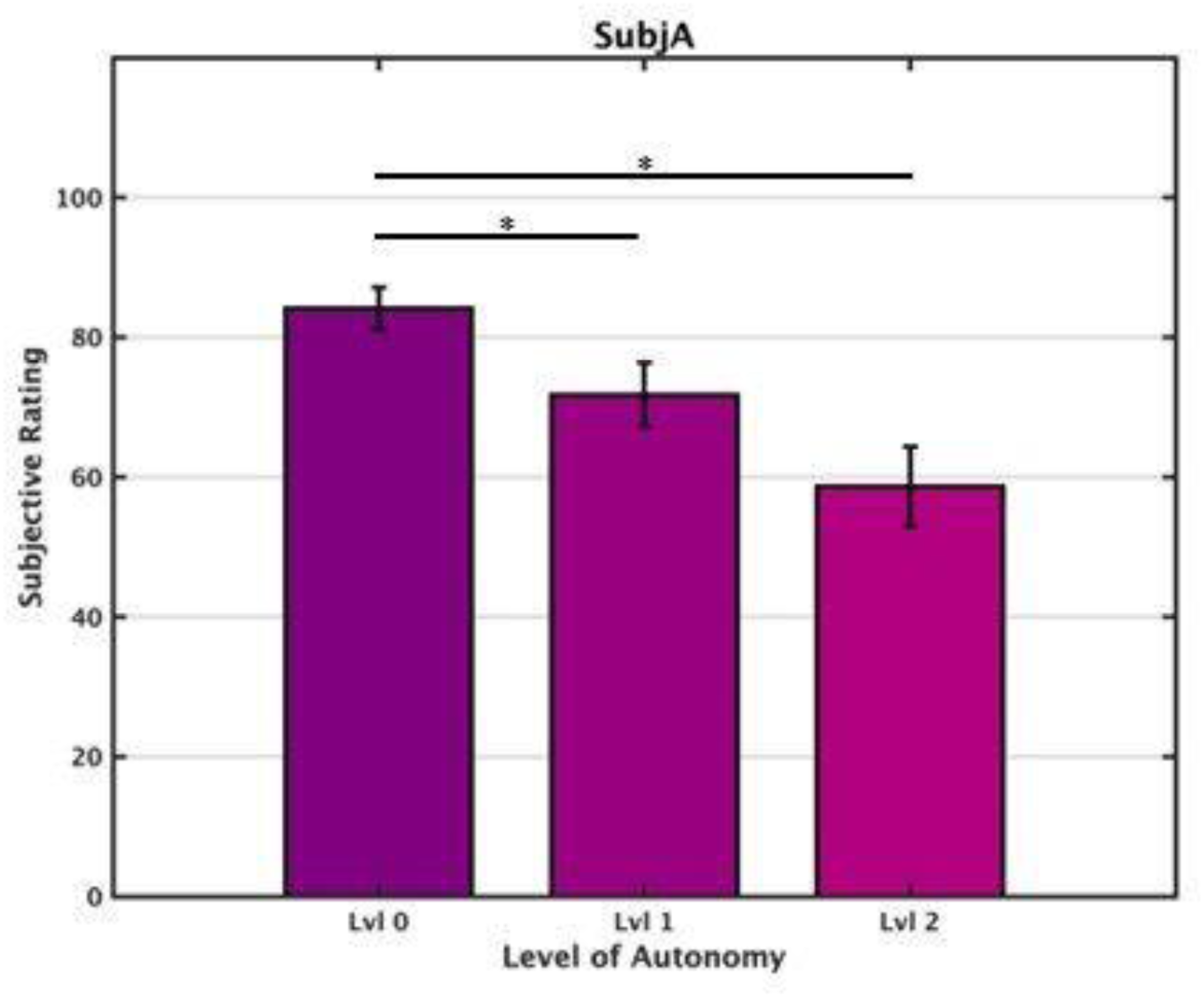
Subjective assessment of responsibility. A significant decrease in subjective judgement of responsibility was found in *Level 0* compared to *Level 1* (p = .006) and in *Level 0* compared to *Level 2* (p <.001), with greater subjective judgement of responsibility in *Level 0* compared to *Level 1* and *Level 2*. * = significant.

## 4. Discussion

In the present pilot study, we aimed to develop a new paradigm to investigate how the sense of agency and (moral) decision-making are influenced by the type of input received from an intelligent autonomous system. To this end, we programmed a task in which participants, in the role of drone operators, had to decide whether (or not) to initiate an attack on a simulated battlefield, during different trial types and with the support of an intelligent system at three levels of autonomy. Ultimately, the overall goal of this research agenda was to develop a paradigm to investigate the mechanisms involved in human-AI interactions in the context of morally challenging situations and to better understand the determining factors in the effect of (inputs received from) autonomous systems on the decisions of human agents faced with moral choices.

Our results show that our new paradigm was sensitive enough to discriminate between moral and non-moral situations. Indeed, participants in our study had a .54 likelihood to initiate an attack in our moral situations, while they refrained from attacking when there were no enemies, and systematically attacked in situations where an enemy was present, but an attack posed no risk for allied troops. Furthermore, the moral situations were characterized by less utilitarian choices than our control condition, reflecting the uncertainty of the situation. Notably, the assistance of the autonomous system increased the number of utilitarian choices, demonstrating the influence of the machine on decision-making. Finally, the moral situations were characterized by a longer reaction time, indicating the participants’ hesitation when they had to decide to fire or to not fire when allies’ life were at stake. Thus, this pilot study thus provides a preliminary paradigm for investigating research questions related to the effects of human-machine interactions in moral decision-making situations.

Our results were partially consistent with our expectations, as the number of attacks increased when there was no risk and decreased when there was no enemy, confirming that our control trials were working properly as well. However, no effects related to the level of autonomy were found regarding the proportion of attacks. One possibility is that human choices, when made in the context of moral dilemmas, are not influenced by the level of autonomy of a system, contrary to choices made in a context without these dilemmas. However, this would be surprising, given that previous studies conducted in military scenarios (e.g., Chen & Joyner, 2009; Rovira et al., 2007) have found an influence of the level of autonomy on human performance and decision. Another interpretation for the lack of an effect of level of autonomy on the rate of A1 choices relies on the way the task was designed (and in particular the possibility for participants to calculate maybe too easily the expected losses for both alternatives in each choice), resulting in ceiling/floor effects in the control conditions and an A1 rate of around 50% during *Moral Decision-Making* trials.

With respect to the Utilitarian Choice, i.e., the proportion of choices that lead to the least loss of allies, we expected an increase in UC in trials without moral conflict and an effect of autonomy leading to an increase in these choices at the highest level of Autonomy (as evidence of an effect of autonomous system on moral decision). Our results confirmed our expectations, with utilitarian choices increasing as a function of Uncertainty and Autonomy. Considering that the recommendations made with the highest level of system support were based on the lower expected losses computed, our results showed that our participants’ moral choices were significantly changed by the input received from the system. This suggests that human choices can be influenced by the recommendations received from a decision support system, independently of the morally-unmorally challenging nature of the situation.

In relation to the Response Time, we expected that participants would take a longer time to make a decision on trials with a moral conflict and Autonomy would shorten participants’ response times. Our results confirmed our expectation, with RTs being longer in moral situations, suggesting that participants took longer to make a decision when a moral conflict was present, and that the highest level of system support shortened their response time in the No-Enemy trials. This last result is consistent with previous findings from laboratory experiments showing that autonomous systems can help users in detection tasks (e.g., Goh et al., 2005), although it is surprising that this effect was found only for the absence of target but not for the presence of target during trials without risk, which is somehow inconsistent with previous findings (e.g., Chavaillaz et al., 2018). One possible reason for this result could be that the target used in our task (i.e., a white square among grey squares on a black background) was too noticeable and thus too easy to recognize for the *Level 1* and *Level 2* functions to be useful to participants.

Regarding SoA, given that a decrease in agency in human-machine interactions has been previously reported (Berberian et al., 2012; Vantrepotte et al., 2022), we firstly expected that participants would show a decrease in the Intentional Binding and subjective sense of responsibility at higher levels of support, indicating a decrease in the SoA. Consistent with our hypothesis, our results showed a decrease in the SoA at the explicit level with higher levels of support. This result is also consistent with the recently described human tendency to attribute moral responsibility to non-human agents which may lead people to be willing to blame them (Furlough, et al., 2021; Kneer and Stuart, 2021; Liu and Du, 2022). Nevertheless, our results did not show a decrease in SoA at the implicit level, which is inconsistent with our expectations and previous studies (Berberian et al., 2012). However, this discrepancy is not completely surprising considering that a dissociation between the two levels of measures in the SoA has already been reported in previous studies (e.g., Synofzik et al., 2008; Moore and Obhi, 2012; Saito et al., 2015). It has been suggested that at the explicit level a higher-order conceptual judgement of being an agent is formed and that this aspect of SoA is closely related to higher-level sources of information such as social and contextual cues (Synofzik et al., 2008), suggesting that Intentional Binding and explicit judgments of agency do not share the same processes (Dewey & Knoblich, 2014). Since the two measurement systems are separable (Saito et al., 2015), it is possible that dissociation between implicit and explicit measurements occurred in our study. In particular, in our new paradigm the three autonomy conditions may not have been so different as to yield a significant difference at the implicit level of agency, but only at the explicit level.

Still regarding SoA, consistent with previous findings (Moretto et al., 2011), we also expected an increase in SoA during Moral Decision-Making trials. Consequently, we expected that moral decision making would also be affected. Because SoA appears to be closely related to moral responsibility (Moretto et al., 2011; Caspar, et al., 2016) and this is reduced by the level of autonomy of the machine, we expected that a decrease in the sense of responsibility would lead to a change in the number of attacks at the highest level of system autonomy. However, contrary to our expectations, the results showed a significant decrease in the SoA in the *Moral Decision-Making* trials at the implicit level (i.e., in the IB). One possible explanation for these results is related to the human tendency to take more responsibility for positive than for negative events, which seems to be a mechanism for increasing self-esteem (Bradley, 1978; Greenberg et al., 1992; Yoshie and Haggard, 2013). However, it has been pointed out there is a tendency to overestimate one’s agency, and that this bias is stronger when the outcome of an action is positive rather than neutral or negative (Wegner & Wheatley, 1999; Haggard, 2017). Because the risk of a potential hit to allies was constantly present in the Moral Decision-Making trials in our task, it is possible that participants in these trials had a reduced SoA and responsibility, and disengaged from the situation due to the risk of negative dramatic consequences of their actions. Alternatively, it is also possible that these results are related to the young cadets’ lower SoA, which has already been described by Caspar and colleagues (Caspar et al., 2018).However, these results should be taken with caution, considering a flaw in the preparation of the Matlab script used to run the experiment, and the possibility that the time intervals used for the time estimations (200, 500, and 800ms) were not completely assigned equally across Uncertainty conditions (i.e., that each time interval are shown five times per Uncertainty condition). Indeed, the program was designed to generate for each three blocks (corresponding to the three levels of Autonomy) 15 presentations of each interval duration (45 intervals in total) presented in a random order across trials (45 trials/block). Albeit randomly presented across Uncertainty conditions, it is possible that the number of presentations of each interval duration was not perfectly the same across Uncertainty condition. Future investigations will be therefore necessary to determine whether the SoA decreases when human subjects interact with autonomous systems when making moral decisions compared to situations that do not pose a moral challenge, or whether this result is due to the methodological flaw observed in our experiment.

To summarize, our results show that human choices made in morally-challenging scenarios can be differentially influenced by the recommendations received from different level of autonomous systems, replicating and extending previous findings (e.g., Chen & Joyner, 2009; Rovira et al., 2007 for studies in the military domain). This was measured by the small but significant difference in the number of utilitarian choices between conditions and the decrement in response time in the No Enemy trials. Interestingly, this effect was completed by a decrement in the subjective measure of agency with higher levels of autonomy, which is consistent with previous research (Barberian et al., 2012; Vantrepotte et al., 2022). At the same time, however, our results show some inconsistencies with previous studies (Moretto et al., 2011; Berberian et al., 2012), with less agency measured in the moral decision-Making trials and no significant difference across Level of Autonomy at the implicit level of agency (IB).Thus, further experiments need to be conducted to determine if the inconsistent results we found were related to the design of the experiment as we suggested above. In particular, in addition to the change need on the Intentional Binding measure, the scenario we used was closer to impersonal/neutral stimuli rather than a moral/emotional context. For example, the radar screen shown to participants was quite schematic and likely did not allow participants to properly imagine the context of the choices they were making. In addition, they did not know the number of victims following their decision, and the victims were not clearly shown as individuals. Thus, it could be that the images we used did not have enough emotional content to reinforce/enhance the SoA. Since the sense of agency could be also affected by the actions’ outcome, which is missing in the current version of the task, to overcome this issue, future research could improve our paradigm by using a less neutral task and content with more moral and emotional valence.

## 5. Conclusion

Considering the increasing presence of intelligent autonomous systems in our daily lives, it is crucial to conduct further research to better understand the implications in sensitive domains such as the military context to provide input for the successful design of innovative automated systems. Our findings suggest that the level of system autonomy influences participants’ moral decision-making and that input received from an intelligent autonomous system influences SoA. By developing a valid paradigm for assessing the impact of human-machine interaction on moral decision-making in the military, with the present study we pave the way for further lines of research on the influence of autonomous systems on human moral behavior, considering the current lack of research on this issue.

## Author contributions

**A.P., S.L.B., E.C.**: Conceptualization, Methodology. **A.P.**: Software. **A.S., A.P.**: Investigation. **A.S., A.P.**: Data curation, Formal Analysis, Writing-Original draft preparation. **E.C., S.L.B.**: Supervision. **S.L.B.**: Funding acquisition. **A.S., A.P., E.C., S.L.B.**: Writing-Reviewing and Editing.

## Funding

This research was funded by Belgian Defense – Royal Higher Institute of Defense, grant number HFM20-03.

## Footnotes

1 This range is commonlyused in human contingency assessment (for recent examples, see Prével et al., 2021 or Vaghi et al., 2019).

## Notes

### Competing Interest Statement

The authors have declared no competing interest.

